# Genetically encoded green fluorescent biosensors for monitoring UDP-GlcNAc in live cells

**DOI:** 10.1101/2021.06.23.449603

**Authors:** Zefan Li, Jing Zhang, Hui-wang Ai

**Affiliations:** Department of Molecular Physiology and Biological Physics, and Center for Membrane and Cell Physiology, University of Virginia, Charlottesville, Virginia 22908, USA

## Abstract

Uridine diphosphate *N*-acetylglucosamine (UDP-GlcNAc) is a nucleotide sugar used by glycosyltransferases to synthesize glycoproteins, glycosaminoglycans, glycolipids, and glycoRNA. UDP-GlcNAc also serves as the donor substrate for the formation of *O*-GlcNAc, a dynamic intracellular protein modification involved in diverse signaling and disease processes. UDP-GlcNAc is thus a central metabolite connecting nutrition, metabolism, signaling, and disease. There is a great interest in monitoring UDP-GlcNAc in biological systems. Here, we present the first genetically encoded, green fluorescent <u>U</u>DP-<u>G</u>lcN<u>Ac</u> <u>s</u>ensor (UGAcS), an optimized insertion of a circularly permuted green fluorescent protein (cpGFP) into an inactive mutant of an *E. coli* UDP-GlcNAc transferase, for ratiometric monitoring of UDP-GlcNAc dynamics in live mammalian cells. Although UGAcS responds to UDP-GlcNAc quite selectively among various nucleotide sugars, UDP and UTP interfere with the response. We thus developed another biosensor named UXPS, which is responsive to UDP and UTP but not UDP-GlcNAc. We demonstrated the use of the biosensors to follow UDP-GlcNAc levels in cultured mammalian cells perturbed with nutritional changes, pharmacological inhibition, and knockdown or overexpression of key enzymes in the UDP-GlcNAc synthesis pathway. We further utilized the biosensors to monitor UDP-GlcNAc concentrations in pancreatic MIN6 β-cells under various culture conditions.

## INTRODUCTION

UDP-GlcNAc, the major end-product of the hexosamine biosynthetic pathway (HBP), is one of the most important nucleotide sugars in living organisms.^1^ The HBP branches out from glycolysis and consumes ~0.006%-3% of total glucose,^2–4^ along with glutamine, acetyl-coenzyme A (Ac-CoA), adenosine triphosphate (ATP), and uridine triphosphate (UTP). Because multiple types of metabolic molecules, including carbohydrates, amino acids, fatty acids, and nucleotides, regulate the flux of HBP, UDP-GlcNAc has been considered an integrator of nutritional and metabolic signals.^5,6^

As an activated *N*-acetylglucosamine (GlcNAc) donor, UDP-GlcNAc is essential for the glycosyltransferase-catalyzed formation of various glycosaminoglycans, glycoproteins, glycolipids, and glycoRNAs.^1,7^ In mammalian cells, glycosylation primarily occurs in the endoplasmic reticulum (ER) and Golgi. The glycosylated products are typically translocated to the extracellular space, playing critical roles such as maintaining structural stability, modulating cell-matrix or cell-cell interaction, regulating cell proliferation and migration, and initiating other types of signaling.^1,6^ Moreover, UDP-GlcNAc is an essential and the only substrate for *O*-GlcNAcylation, a reversible post-translational modification of nucleocytoplasmic proteins.^2,8,9^ *O*-GlcNAc transferase (OGT) catalyzes the transfer of the GlcNAc subunit from UDP-GlcNAc to the serine or threonine residues of proteins, while O-GlcNAcase (OGA) hydrolyzes the modification to generate free proteins and GlcNAc. This dynamic and tightly regulated *O*-GlcNAcylation process, analogous to more well-known phosphorylation, is involved in a large array of intracellular signaling processes.^1,5,6,10–12^ Aberrant *O*-GlcNAcylation has been linked to aging, neurodegeneration, cancer, cardiovascular diseases, and metabolic disorders.^2,6,13–15^

The concentration of UDP-GlcNAc is one of the several key factors regulating glycosylation. The drastic increase of the β1,6-branched oligosaccharide levels was observed in B16 melanoma cells incubated with GlcNAc.^16^ In another example, deleterious mutations in SLC35A3, the major Golgi UDP-GlcNAc transporter, were identified in patients with autism spectrum disorder, arthrogryposis, and epilepsy.^17^ These mutations reduce UDP-GlcNAc transport into the ER, leading to a massive decrease of highly branched *N*-glycans and a drastic increase of lower branched glycoforms at the cell surface.^17^ The UDP-GlcNAc level has also been found to regulate intracellular *O*-GlcNAcylation. *In vitro* characterization of OGT demonstrated that the concentration of UDP-GlcNAc was positively correlated with the *O*-GlcNAcylation of the tested peptide substrates.^18^ Supplementing human hepatocellular carcinoma HepG2 cells with glucosamine, a metabolite used by HBP to synthesize UDP-GlcNAc, significantly increased *O*-GlcNAc.^19,20^ Furthermore, hyperglycemia was shown to increase the *O*-GlcNAc level in multiple tissue types via the HBP. ^21–23^

Because of the importance of UDP-GlcNAc in metabolic sensing, signaling, and disease, methods for monitoring UDP-GlcNAc levels in living systems are highly needed. Traditionally, chromatography methods are used to determine cellular UDP-GlcNAc levels,^3,24,25^ but these methods require cell lysis and provide little spatiotemporal resolution. To address this technical gap, we engineered the first genetically encoded fluorescent sensor, UGAcS, for detecting UDP-GlcNAc in living cells. We inserted a cpGFP into an inactive mutant of murG,^26–28^ an *E. coli* UDP-GlcNAc transferase, and performed directed evolution to optimize the biosensor. Because UDP and UTP interfere with the response of UGAcS to UDP-GlcNAc, we developed an additional control biosensor, UXPS, which is only responsive to UDP and UDP. We demonstrated the use of the biosensors to follow UDP-GlcNAc concentration changes in cultured mammalian cells in response to various nutritional, pharmacological, and genetic perturbations.

## RESULTS AND DISCUSSION

### Design, Engineering, and in vitro characterization of the UGAcS

MurG is a well-characterized *E. coli* UDP-GlcNAc transferase involved in synthesizing lipid-linked precursors to assemble peptidoglycan, the polymeric cell wall outside the bacterial cell membrane.^28,29^ MurG has a high binding affinity to UDP-GlcNAc (~1.4 μM), and its structures in the apo and UDP-GlcNAc-bound forms have been reported.^27,29^ We selected murG as the sensory domain to build a UDP-GlcNAc sensor. By carefully examining the structures of murG, we identified that the binding of UDP-GlcNAc triggers a structural conversion of residues 60-70 in MurG from a loop into an α-helix (**Supplementary Information, Figure S1**). We further confirmed the significant conformation change at this loop by analyzing the changes of dihedral angles of every four consecutive C_◻_ atoms (**Figure S2**).

We next inserted cpGFP to the above-identified loop between residues 64 and 65 of murG (**Figure 1A** & **Figure S3**). A fully randomized residue was introduced to each of the two junctions as the linkers. We screened the library and identified a UGAcS0.1 mutant with a 30% response (ΔR/R_0_ or (R-R_0_)/R_0,_ where R is the ratio of fluorescence with 488 nm excitation to that with 400 nm excitation) to UDP-GlcNAc. To increase the sensor’s response to UDP-GlcNAc, we performed three rounds of error-prone PCR, and the screening of these libraries resulted in UGAcS0.2 showing a 300% (ΔR/R_0_) response.

**Figure 1.**
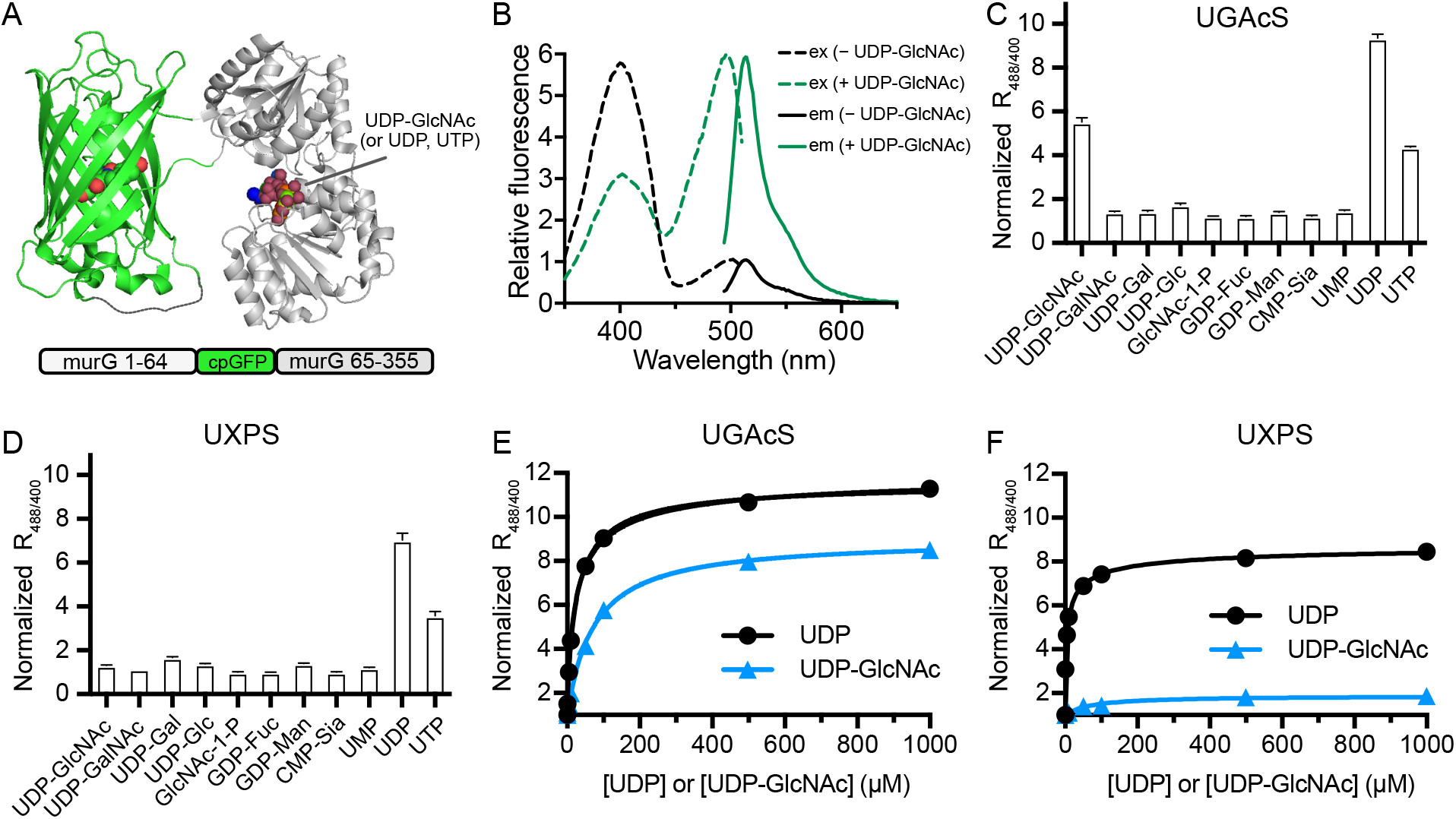
Design and *in vitro* characterization of the biosensors. (**A**) Illustration of the sensor design, showing the cpGFP insertion between residues 64 and 65 of the murG glycosyltransferase. Also highlighted is a UDP-GlcNAc molecule in the substrate-binding pocket. (**B**) Fluorescence excitation and emission spectra of UGAcS before and after addition of 1 mM UDP-GlcNAc. (**C**, **D**) Responses of UGAcS (C) or UXPS (D), presented as normalized fluorescence excitation ratios (488 nm / 400 nm), to 100 μM of various nucleoside sugars and other related cellular metabolites. Data are presented as mean ± SD (n=3). (**E**, **F**) Dose-dependent responses of UGAcS (E) or UXPS (F) to UDP or UDP-GlcNAc.

Although our primary goal is to apply the fluorescent biosensors in mammalian cells and murG is unlikely to be active in mammalian cells where its lipid-linked peptidoglycan substrate is not present, we still performed saturation mutagenesis on the catalytic His19 residue of MurG in UGAcS0.2. This residue is essential for catalysis and conserved in murG from 73 orthologues.^26,27^ Our screening of the mutants led to an enzymatically inactive His19Ser mutant (UGAcS0.3) but comparable to UGAcS0.2 in terms of the UDP-GlcNAc responsiveness.

Because UDP is a natural inhibitor and regulator of UDP-sugar transferases, we examined the fluorescence of UGAcS0.3 upon the addition of UDP. UGAcS0.3 showed a higher response to UDP than UDP-GlcNAc. We next devoted our effort to engineering UGAcS0.3 for increased specificity to UDP-GlcNAc versus UDP. We chose five pairs of residues (residues 192 and 193, residues 16 and 127, residues 164 and 269, or residues 244 and 245) in the ligand-binding pocket and performed saturation mutagenesis. Screening of these libraries resulted in UGAcS0.4 with mutations at residues 192 and 193. In comparison with UGAcS0.3, UGAcS0.4 showed increased responsiveness to UDP-GlcNAc and reduced responsiveness to UDP. Despite the progress, we were unable to identify a mutant to exclude the UDP interference entirely.

From UGAcS0.4, we performed two more rounds of random mutagenesis on UGAcS0.4. By screening these libraries for improved UDP-GlcNAc responsiveness, we arrived at UGAcS, which showed nearly 700% response (ΔR/R_0_) to 1 mM UDP-GlcNAc (**Figure 1B** & **Figure S4**). We further tested the specificity of UGAcS using various nucleotide sugars and other related compounds at physiologically relevant concentrations (**Figure 1C**). UGAcS responded to UDP-GlcNAc, UDP, and UTP. Other tested nucleotide sugars, including uridine diphosphate *N*-acetylgalactosamine (UDP-GalNAc) and uridine diphosphate glucose (UDP-Glc) that are structurally very close to UDP-GlcNAc, induced little fluorescence change.

After we realized that it might be unrealistic to engineer a fully specific UDP-GlcNAc biosensor from the glycosyltransferase, we sought to identify a UDP sensor, which could be used as a control to cross-check the responses of the UGAcS variants. From one of the above-mentioned ligand-binding-pocket mutagenesis libraries, we identified a mutant (UXPS), which is responsive to UDP and UTP, not to UDP-GlcNAc and other nucleotide sugars (**Figure 1D** & **Figure S4**).

We next titrated UGAcS and UXPS with various concentrations of UDP-GlcNAc and UDP (**Figure 1EF)**. The apparent dissociation constant (*K*_d_) values (deduced from the fluorescence responses of the sensors) for UGAcS were determined to be 72±4 μM and 26±1 μM in the presence of UDP-GlcNAc and UDP, respectively, while UXPS responded to UDP with an apparent *K*_d_ of 6±1 μM.

### Monitoring of UDP-GlcNAc changes in HEK 293T cells

2-Deoxy-D-glucose (2-DG) is a glucose analog, and 2-deoxy-D-glucose-6-phosphate (2-DG6P) intracellularly formed from 2-DG competitively inhibits hexokinase (HK) and phosphoglucose isomerase (PGI), the enzymes involved in the first two steps of glycolysis (**Figure 2A**).^30^ To validate the function of UGAcS in mammalian cells, we expressed the sensors in human embryonic kidney (HEK) 293T cells and examined sensor responses to glycolysis inhibition by 2-DG. HEK293T cells transiently expressing UGAcS emitted strong green fluorescence. Upon 3-hour incubation with 10 mM 2-DG, the ratio of the fluorescence with 488 nm excitation to that with 400 nm excitation (denoted as R_488/400_) decreased by ~ 15.1% (**Figure 2B**). Meanwhile, we expressed the control sensor UXPS in HEK 293T cells and treated the cells using the same procedure. The fluorescence change of UXPS seemed to be in the opposite direction to that of UGAcS, although the magnitude of the difference was not statistically significant (**Figure 2C**).

**Figure 2.**
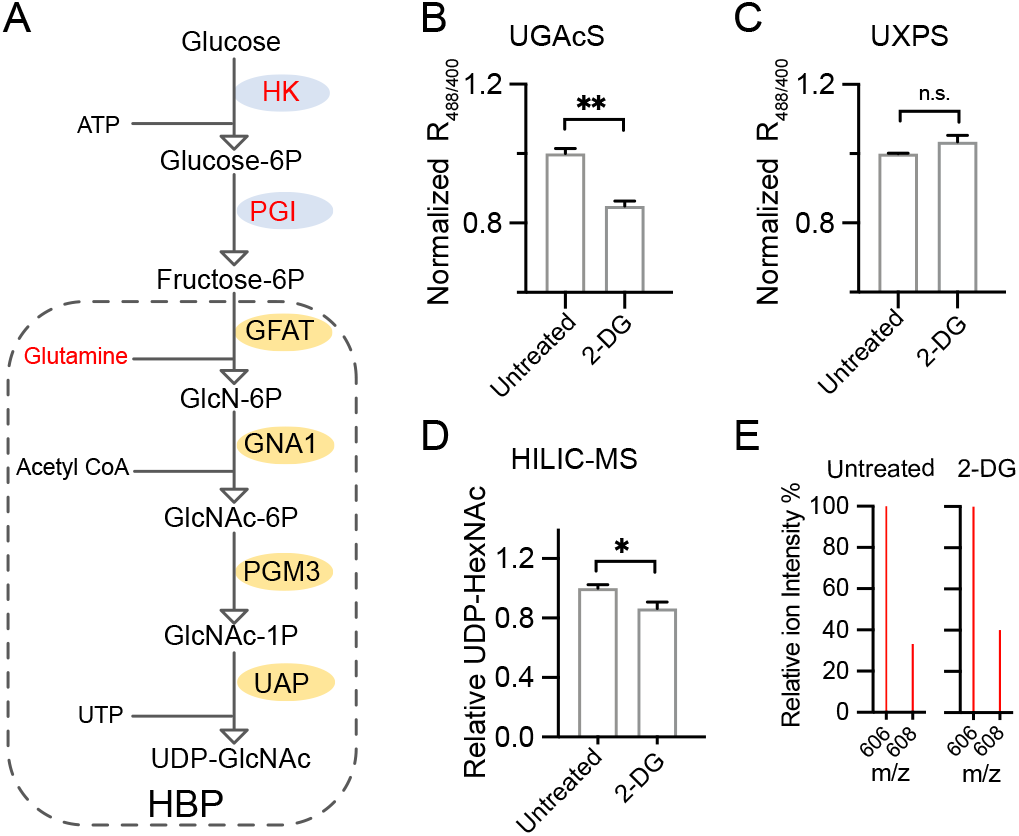
UDP-GlcNAc level changes in HEK 293T cells perturbed with 2-DG. (**A**) Schematic illustration of the hexosamine biosynthetic pathway (HBP) responsible for the production of UDP-GlcNAc. 2-DG (2-deoxy-D-glucose) is a glycolysis inhibitor acting on hexokinase (HK) and phosphoglucose isomerase (PGI). (**B**, **C**) Responses of UGAcS (B) or UXPS (C), given as normalized fluorescence excitation ratios (488 nm / 400 nm), to 3-hour incubation with 10 mM 2-DG. (**D**) HILIC-MS analysis of relative UDP-HexNAc concentrations in extracts of HEK 293T cells untreated or treated with 10 mM 2-DG. (**E**) Representative mass spectrograms of the samples in panel D. The m/z = 606 peak represents the major isotope peak for UDP-HexNAc. ^13^C-double-labeled UDP-GlcNAc (^13^C_2_-UDP-GlcNAc) was doped into the cell extracts as an internal standard, contributing primarily to the m/z = 608 peak. Data in panels B-D are presented as mean ± SEM (n=3). *P* values were determined by one-way ANOVA with Dunnett’s multiple comparisons test (\*\**P*<0.01; *P<0.05; and n.s., not significant, *P*≥0.05).

To cross-verify the results, we adapted a hydrophilic interaction chromatography-mass spectrometry (HILIC-MS) method to quantify UDP-GlcNAc from cell extracts..^3,31^ Briefly, we prepared the lysates of HEK 293T cells treated or untreated with 2-DG, and dopped in ^13^C-double-labeled UDP-GlcNAc (^13^C_2_-UDP-GlcNAc) as an internal standard. The samples were separated on a zwitterionic polymer-based high performance liquid chromatography (HPLC) column before being fused into an electrospray ionization (ESI) single quadrupole mass spectrometer. 606 and 608 m/z ions, which are the major isotope peaks for UDP-GlcNAc and ^13^C_2_-UDP-GlcNAc, respectively, were monitored (**Figure S5**). The intensity ratio of the two peaks is thus an indicator for the relative UDP-GlcNAc concentrations in the cell lysates. Since our chromatographic condition did not separate UDP-GlcNAc from UDP-GalNAc and the two nucleotide sugars have identical molecular formulas, the HILIC-MS method in fact measured UDP-GlcNAc and UDP-GalNAc (referred to as UDP-HexNAc) collectively. Specific epimerases are responsible for the interconversion of UDP-GlcNAc and UDP-GalNAc in cells. The concentration ratio of UDP-GlcNAc to UDP-GalNAc is usually ~ 3:1.^32^ Thus, the UDP-HexNAc measurements from HILIC-MS can be used to approximate UDP-GlcNAc level changes. The HILIC-MS analysis confirmed that 2-DG reduced the UDP-HexNAc level by ~ 13.4% (**Figure 2DE**), and the result is very well aligned with the UGAcS-based fluorescence assay.

The glutamine fructose-6-phosphate aminotransferase (GFAT) is a feedback-regulated rate-limiting enzyme in the HBP.^33^ Since glucosamine (GlcN) enters the HBP downstream of GFAT (**Figure 2A**), GlcN is a potent stimulator of HBP and can rapidly increase the intracellular concentration of UDP-GlcNAc.^20^ We used a confocal microscope equipped with 488 and 405 nm lasers to follow GlcN-stimulated UDP-GlcNAc elevation in HEK 293T cells. Upon GlcN stimulation, the fluorescence with 488 nm excitation increased along with the simultaneous decrease of the fluorescence with 405 nm excitation (**Figure 3AB**), resulting in an overall ~ 70% ratiometric change (ΔR/R_0_). Most of the change was completed within the first 30 min post-stimulation. In contrast, GlcN triggered an ~ 33% ratiometric change (ΔR/R_0_) of the UXPS fluorescence in the opposite direction (**Figure 3CD)**. The observed fluorescence change of UXPS is not surprising because the GlcN-dependent UDP-GlcNAc synthesis process may consume UTP quickly. Although the opposite responses of UGAcS and UXPS can exclude the possibility of the observed UGAcS response being caused by pH changes, we still further tested cpGFP-expressing HEK 293T cells against GlcN stimulation and observed no response (**Figure 3EF)**. Collectively, these results confirmed that GlcN indeed increased the intracellular UDP-GlcNAc concentration and that UGAcS successfully detected this relatively rapid increase of UDP-GlcNAc in HEK 293T cells.

**Figure 3.**
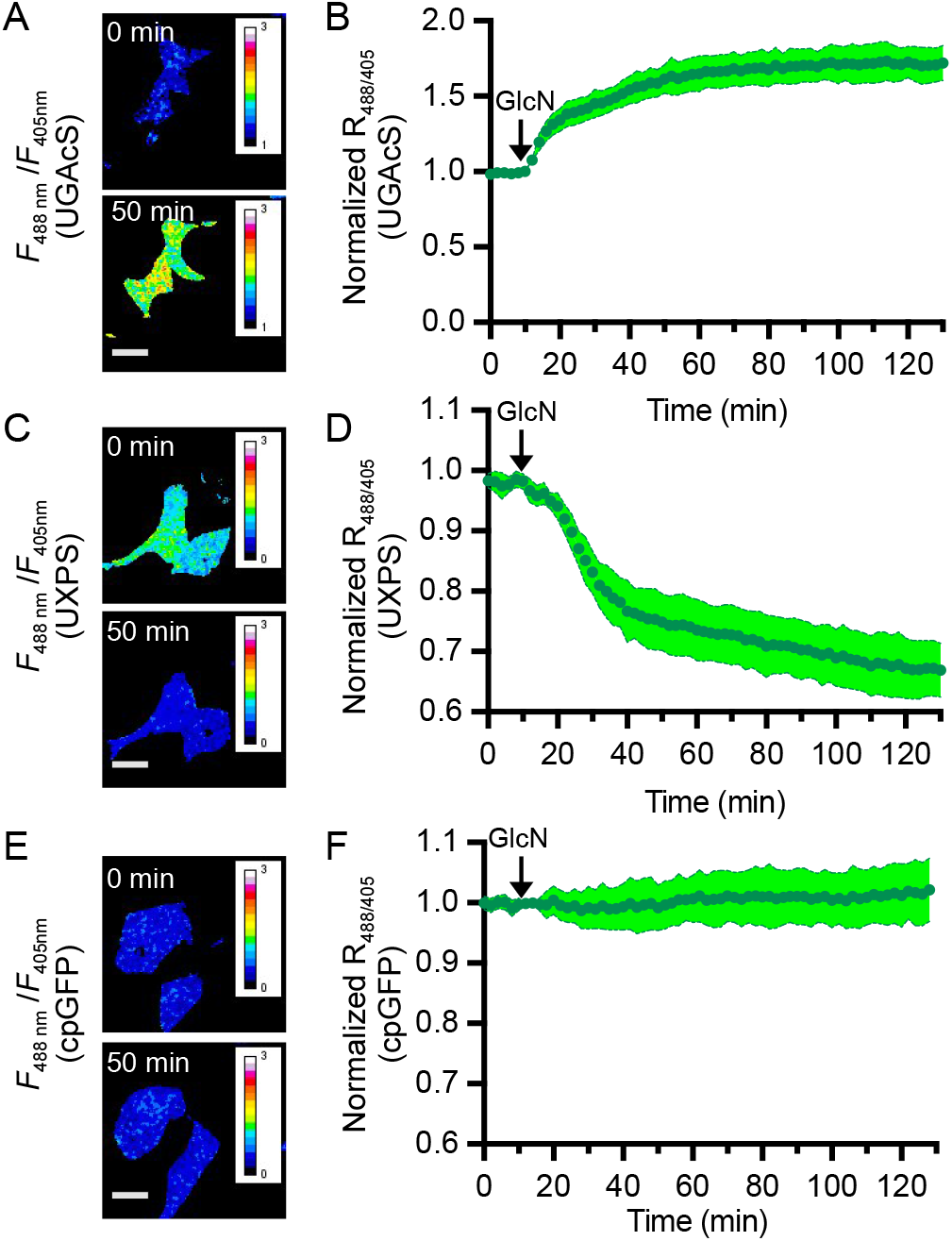
Glucosamine (GlcN)-induced UDP-GlcNAc increase in HEK 293T cells. (**A**, **C, E**) Representative ratiometric images of HEK 293T cell expressing UGAcS (A), UXPS (C), or cpGFP (E) with two excitation wavelengths (488 nm / 405 nm) before and after treatment with 5 mM GlcN (scale bars, 40 μm). (**B, D, F**) Quantitative traces of normalized fluorescence excitation ratios (488 nm / 405 nm) for UGAcS (B), UXPS (D), or cpGFP (F) in HEK 293T cells. Data are presented as mean ± SD (n=10 cells from 3 cultures).

We next used the biosensors to examine UDP-GlcNAc concentration changes in response to the genetic disruption of two key enzymes in the HBP. GFAT is the first and rate-limiting enzyme, while UDP-*N*-acetylglucosamine pyrophosphorylase (UAP) is the last enzyme in the pathway and responsible for the direct synthesis of UDP-GlcNAc (**Figure 4A**). We used short hairpin RNAs (shRNAs) to knock down GFAT or UAP. The effectiveness of the shRNAs was first confirmed using fluorescence assays with HEK 293T cells co-expressing corresponding shRNAs and the GFAT1 or UAP1 gene fused to a red fluorescent protein (RFP) mScarlet-I via a P2A self-cleaving peptide (**Figure S6**). A scramble non-targeting shRNA sequence (shNC) was used as a negative control, and the GFAT or UAP shRNAs that induced the most significant decrease of mScarlet-I fluorescence were selected for further experiments. Next, we used the shRNA lentiviral vectors to infect HEK 293T cells, to which the gene of UGAcS or UXPS was further introduced by transfection. As expected, GFAT or UAP knockdown decreased the UDP-GlcNAc level, as the fluorescence excitation ratios (R_488/400_) of UGAcS were reduced compared to the shNC control group (**Figure 4B**). Meanwhile, the fluorescence excitation ratios (R_488/400_) of UXPS in the experimental groups were slightly higher (statistically insignificant) than the shNC control group (**Figure 4C**).

**Figure 4.**
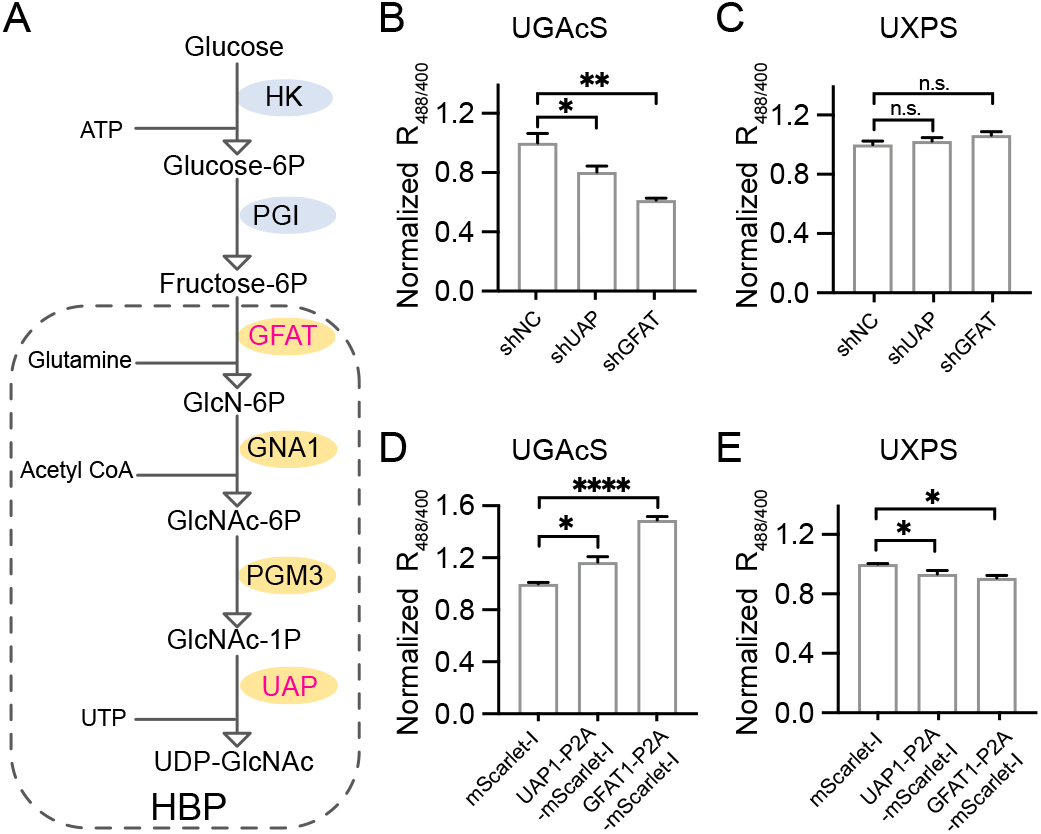
UDP-GlcNAc level changes in response to knockdown or over-expression of GFAT or UAP in HEK 293T cells. (**A**) Schematic illustration of the HBP UDP-GlcNAc synthesis pathway. Highlighted in magenta are the two enzymes selected for genetic manipulation in this study. (**B**, **C**) Responses of UGAcS (B) or UXPS (C), presented as normalized fluorescence excitation ratios (488 nm / 400 nm), to shRNA knockdown of GFAT or UAP. A negative shRNA control, shNC, was used for comparison and normalization. (**D**, **E**) Responses of UGAcS (D) or UXPS (E), given as normalized fluorescence excitation ratios (488 nm / 400 nm), to the overexpression of UAP1-P2A-mScarlet-I or GFAT1-P2A-mScarlet-I. The overexpression of mScarlet-I alone was used for comparison and normalization. Data in panels B-E are presented as mean ± SEM (n=3). *P* values were determined by one-way ANOVA with Dunnett’s multiple comparisons test (\*\*\*\**P*<0.0001; \*\**P*<0.01; *P<0.05; and n.s., not significant, P≥0.05).

We also examined the impact of overexpressing GFAT1 or UAP1 on the UDP-GlcNAc level. The sensor responses are opposite to those in the knockdown experiments: GFAT1 or UAP1 overexpression increased the fluorescence excitation ratios (R_488/400_) of UGAcS and decreased the fluorescence excitation ratios (R_488/400_) of UXPS (**Figure 4DE**). Moreover, in both the knockdown and overexpression experiments, manipulating the GFAT1 expression level induced more dramatical fluorescence responses than manipulating UAP1. This is again expected because GFAT plays the rate-limiting role in the HBP.

### Imaging of UDP-GlcNAc levels in pancreatic MIN6β-cells

Pancreatic β-cells are responsible for the synthesis and secretion of insulin, a key endocrine regulator of glucose levels in the blood and other tissues.^34^ The glucose metabolism of pancreatic β-cells is tightly coupled to insulin synthesis and secretion.^34^ Meanwhile, *O*-GlcNAc levels in β-cells have been linked to the regulation of insulin gene expression, proinsulin-to-insulin processing, and glucose-stimulated insulin secretion.^35–37^ In this context, we used our new biosensors to examine UDP-GlcNAc levels in MIN6 β-cells, a mouse insulinoma cell line, in response to various nutritional conditions. Expression of UGAcS or UXPS in MIN6 cells resulted in bright green fluorescence. Upon 5 mM GlcN stimulation, the fluorescence excitation ratios (R_488/400_) of UGAcS-expressing cells increased by ~ 30% (**Figure 5AB**) within one hour, while ~ 25% decrease was observed for UXPS-expressing cells (**Figure 5CD**). The results suggest that GlcN, which bypasses the rate-limiting GFAT in the HBP, can stimulate the biosynthesis of UDP-GlcNAc in MIN6 cells in a manner similar to that in HEK 293T cells. Next, we cultured UGAcS- or UXPS-expressing MIN6 cells in media with various glucose concentrations (0 mM, 2 mM, and 25 mM) for 20 h (**Figure 5E-H**). The fluorescence excitation ratio (R_488/400_) of UGAcS under the no glucose condition was ~ 50% lower than the 2 mM glucose condition, while we only observed a quite minimal increase (statistically insignificant) of the fluorescence excitation ratio (R_488/400_) of UGAcS from 2 mM to the 25 mM glucose condition. The changes in the fluorescence excitation ratio (R_488/400_) of UXPS were not significant under all conditions. Taken together, the UDP-GlcNAc level in MIN6 cells was sensitive to severe hypoglycemia, but relatively insensitive to glucose concentrations changes from 2 to 25 mM. These findings corroborate a recent study on the mouse heart tissue with 5.5 mM and 25 mM glucose, which concluded that glucose availability alone does not regulate the HBP flux.^3^ In other studies, UDP-GlcNAc has been shown to directly inhibit GFAT via a negative feedback mechanism.^33,38^ The tight regulation of the HBP, however, does not necessarily undermine the importance of UDP-GlcNAc in nutritional signaling. First, multiple types of metabolic molecules are funneled into the HBP, and a collection of these molecules may be needed to drastically shift the HBP flux. In addition, the GFAT expression level and activity are responsive to other signal inputs, and the HBP is thus likely regulated to different extents in diverse tissue types and under various pathophysiological conditions.^39–41^

**Figure 5.**
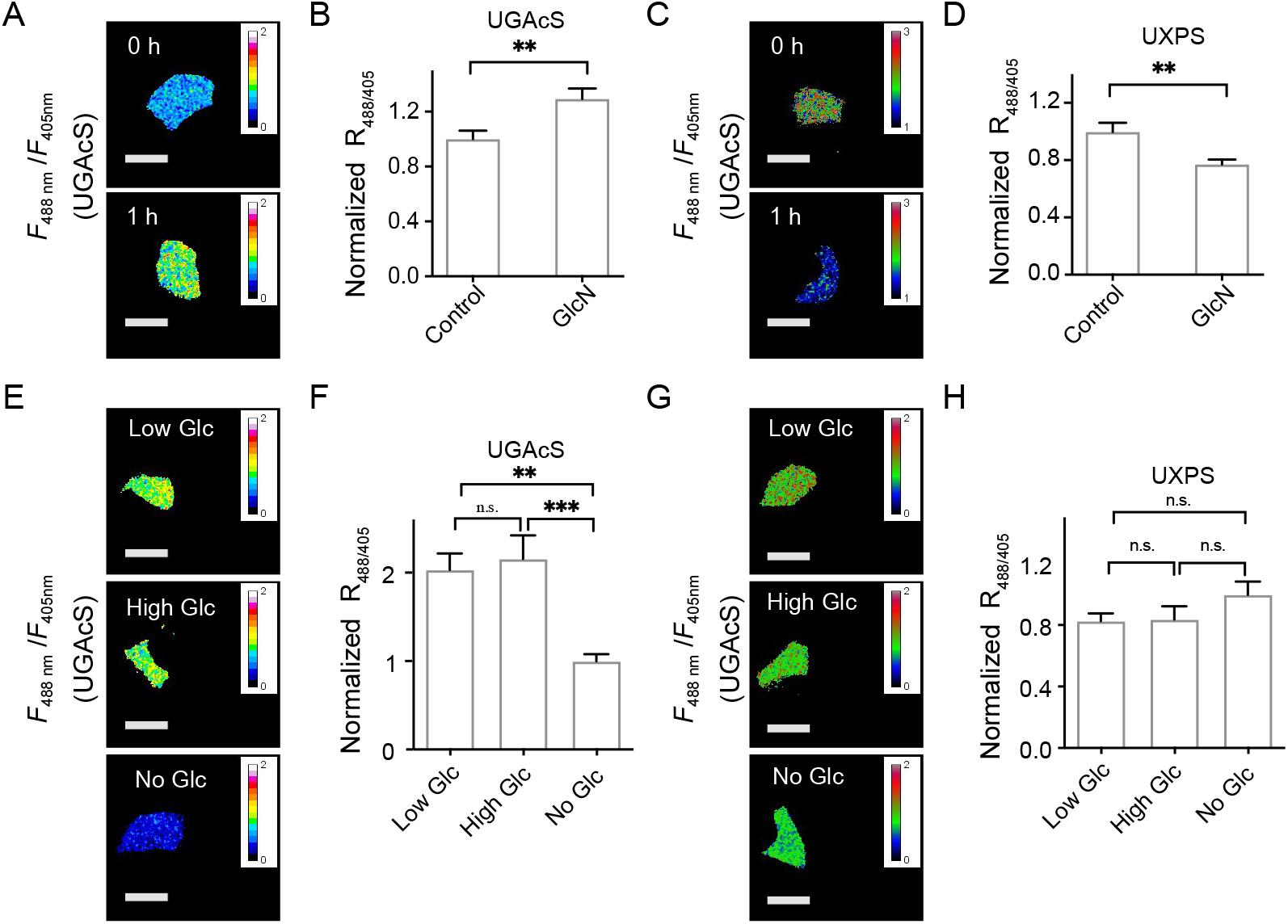
Imaging of UDP-GlcNAc levels in pancreatic MIN6 β-cells. **(A, C)** Representative ratiometric images of MIN6 cell expressing UGAcS (A), UXPS (C) with two excitation wavelengths (488 nm / 405 nm) before and after 1-hour treatment with 5 mM GlcN (scale bars, 40 μm) **(B, D)** Responses of UGAcS (B) or UXPS (D), given as normalized fluorescence excitation ratios (488 nm / 405 nm), to 1-hour incubation with 5 mM GlcN. **(E, G)** Representative ratiometric images of MIN6 cells expressing UGAcS (E), UXPS (G) with two excitation wavelengths (488 nm / 405 nm) cultured in low (2 mM), high (25 mM) or no glucose DMEM for 20 h (scale bars, 40 μm). (**F, H**) Responses of UGAcS (F) or UXPS (H), given as normalized fluorescence excitation ratios (488 nm / 405 nm) to low (2 mM), high (25 mM) or no glucose DMEM for 20 h. Data in panels B, D, F and H are presented as mean ± SEM (n=12). *P* values in panels B and D were determined by unpaired t test (\*\**P*<0.01). *P* values in panels F and H were determined by one-way ANOVA with Dunnett’s multiple comparisons test (\*\*\**P*<0.001; \*\**P*<0.01; and n.s., not significant, *P*≥0.05).

## CONCLUSION

We have engineered a genetically encoded UDP-GlcNAc sensor (UGAcS) by inserting cpGFP into an inactivated UDP-GlcNAc transferase. Because UGAcS is also responsive to UDP and UTP, we further engineered a control sensor, UXPS, which is only responsive to UDP and UTP but not UDP-GlcNAc. We successfully applied the biosensors to monitoring UDP-GlcNAc level changes in HEK 293T cells in response to 2-DG-induced glycolysis inhibition, glucosamine stimulation, and the genetic manipulation of two key enzymes (GFAT and UAP) in the HBP. Finally, we used our biosensors to monitor UDP-GlcNAc levels in pancreatic MIN6 β-cells under various cell culture conditions. Our results suggest that glucose metabolism via the HPB is tightly regulated across a large glucose concentration range. Further research is clearly needed to further understand the regulatory mechanism of the HBP flux, and the fluorescent biosensors described here should facilitate these studies.

In addition to being used as research tools in studying the crucial roles of UDP-GlcNAc in disease and normal conditions, the biosensors may have translational applications. Modulation of the HBP has been considered as a promising method to treat diseases, such as cancer and diabetes.^42,43^ We envision the use of the biosensors to screen for chemical or genetic modulators of the HBP. In addition, murG is a key enzyme for peptidoglycan synthesis in bacteria,^44^ so these murG-based biosensors may be used to discover novel murG inhibitors as a new class of antibiotics.

Our effort to develop the first genetically encoded nucleotide sugar sensor will spur the development of future biosensors. UDP and UTP interfere with the response of UGAcS. Although UXPS can be used for cross-checking, a subset of treatment conditions may move the fluorescence ratios of both sensors toward the same direction, and it would become difficult to interpret the results. In addition, the interference makes the quantitative measurement of intracellular UDP-GlcNAc levels difficult. Directed evolution, machine learning, and computation-assisted design may be combined to further tune the specificity of UGAcS. Also, it may be possible to use alternative strategies to develop fluorescent UDP-GlcNAc sensors with different selectivity profiles. Furthermore, future research may lead to biosensors with altered affinities, in additional fluorescence colors, and for other important nucleotide sugars and carbohydrates.

## Supporting Information

The following Supporting Information is provided: Figures S1−S6, experimental methods, and supplementary references.

## Author Contributions

J.Z. constructed and evaluated the plasmids for shRNA knockdown. Z.L. performed all other experiments, including sensor engineering and characterization, analyzed data, prepared figures, and wrote the manuscript. H.W.A. supervised research, prepared figures, and wrote the manuscript.

## Notes

The authors declare no competing financial interest. The plasmids for bacterial and mammalian expression of UGAcS and UXPS and their sequence information have been deposited to Addgene (Plasmids #172135, #172136, #172137, and #172138).

## Acknowledgments

Research reported in this publication was supported by the NIH Common Fund Glycoscience Program and the National Cancer Institute of the National Institutes of Health under Award Number U01CA230817. The content is solely the responsibility of the authors and does not necessarily represent the official views of the National Institutes of Health. pMD2.G (Addgene Plasmid #12259) and psPAX2 (Addgene Plasmid #12260) were gifts from Didier Trono (EPFL). pLKO.1 (Addgene Plasmid #10878) was a gift from David Root (Broad Institute). We also thank Yu Pang for the independent replication of some results in Figure 1, and Dr. Yiyu Zhang for assisting in culturing MIN6 cells.

