## Supplementary Information for "Genetically encoded green fluorescent biosensors for monitoring UDP-GlcNAc in live cells"

### SUPPLEMENTARY FIGURES

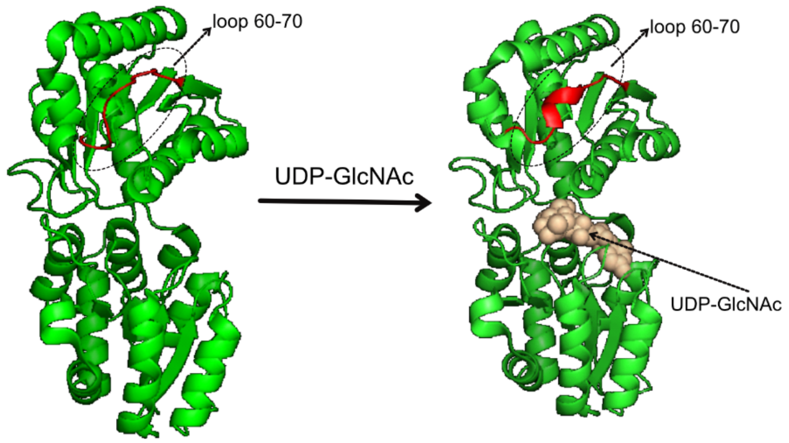


**Figure S1.** Structures of murG before (PDB 1f0k) and after (PDB 1nlm) binding the UDP-GlcNAc ligand, showing the ligand-induced structural change of residues 60-70 (highlighted in red) from a free loop to an α-helix.

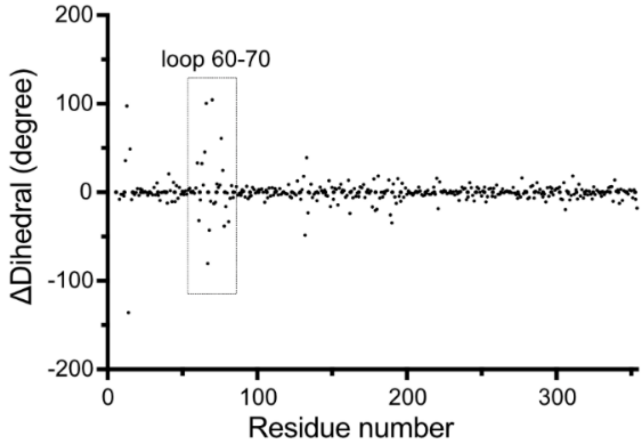


**Figure S2.** $C_\alpha$ dihedral differences (based on every four consecutive $C_\alpha$ atoms) between the ligand-bound (PDB 1nlm) and apo (PDB 1f0k) structures of murG plotted against murG residue numbers.

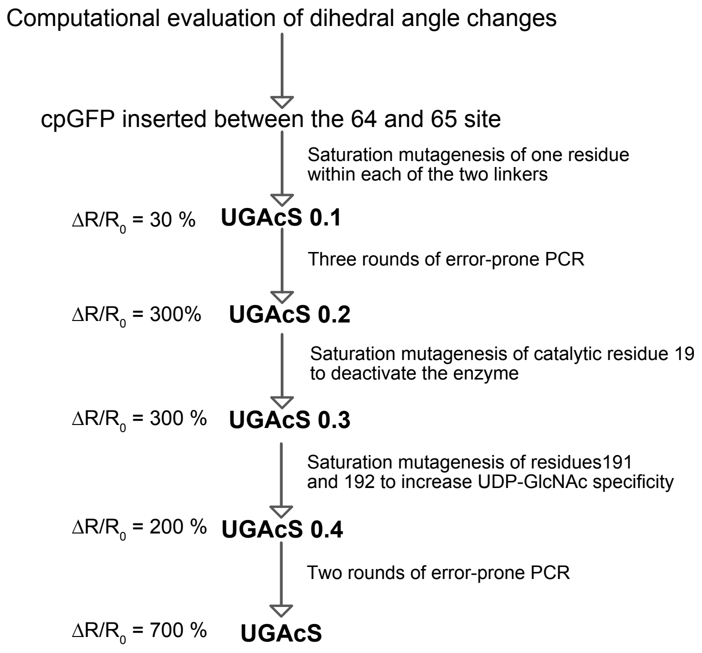


**Figure S3.** Flowchart to illustrate our multi-step process to engineer UGAcS. $\Delta R/R_0$ for each sensor generation is also presented.

```
UGAcS        MSGQGKRLMVIAGGTGGSVFPGLAVAHHLMAQGWQVRWLGTADRMEADLVPKHGIEIDFI
UXPS         MSGQGKRLMVMAGGLGGSVFPGLAVAHHLMAQGWQVRWLGTADRMEADLVPKHGIEIDFI
UGAcS0.1     MSGQGKRLMVMAGGTGGHVFPGLAVAHHLMAQGWQVRWLGTADRMEADLVPKHGIEIDFI
UGAcS0.2     MSGQGKRLMVMAGGTGGHVFPGLAVAHHLMAQGWQVRWLGTADRMEADLVPKHGIEIDFI

UGAcS        RISGEVYIKADKQKNGIKANFQIRHNVEDGSMQLADHYQQNTPIGDGPVLLPDNHYLSTQ
UXPS         RISGEVYIKADKQKNGIKANFQIRHNVEDGSMQLADHYQQNTPIGDGPVLLPDNHYLSTQ
UGAcS0.1     RISGRVYIKADKQKNGIKANFQIRHNVEDGSVQLADHYQQNTPIGDGPVLLPDNHYLSHQ
UGAcS0.2     RISGEVYIKADKQKNGIKANFQIRHNVEDGSMQLADHYQQNTPIGDGPVLLPDNHYLSTQ

UGAcS        SVLSKDPNEKRDHMVLLEFVTAAGITLGMDELHKVDGGSGGTGVSRGEELFTGVVPILVE
UXPS         SVLSKDPNEKRDHMVLLEFVTAAGITLGMDELHKVDGGSGGTGVSRGEELFTGVVPILVE
UGAcS0.1     SVLSKDPNEKRDHMVLLEFVTAAGITLGMDELHKVDGGSGGTGVSKGEELFTGVVPILVE
UGAcS0.2     SVLSKDPNEKRDHMVLLEFVTAAGITLGMDELHKVDGGSGGTGVSRGEELFTGVVPILVE

UGAcS        LDGDVNGHKFRVRGEGEGDATNGKLTLKFICTTGKLPVPWPTLVTTLTYGVQCFSRYPDH
UXPS         LDGDVNGHKFRVRGEGEGDATNGKLTLKFICTTGKLPVPWPTLVTTLTYGVQCFSRYPDH
UGAcS0.1     LDGDVNGHKFRVRGEGEGDATNGKLTLKFICTTGKLPVPWPTLVTTLTYGVQCFSRYPDH
UGAcS0.2     LDGDVNGHKFRVRGEGEGDATNGKLTLKFICTTGKLPVPWPTLVTTLTYGVQCFSRYPDH

UGAcS        MKQHDFFKSAMPEGYVQERTIFFKDDGTYKTRAEVKFEGDTLVNRIELKGIDFKEDGNIL
UXPS         MKQHDFFKSAMPEGYVQERTIFFKDDGTYKTRAEVKFKGDTLVNRIELKGIDFKEDGNIL
UGAcS0.1     MKQHDFFKSAMPEGYVQERTIFFKDDGTYKTRAEVKFEGDTLVNRIELKGIDFKEDGNIL
UGAcS0.2     MKQHDFFKSAMPEGYVQERTIFFKDDGTYKTRAEVKFEGDTLVNRIELKGIDFKEDGNIL

UGAcS        GHKLEYESLRGKGIKALIAAPLRIFNAWEQARAIMKAYKPDVVLGMGGYVSGPGGLAAWS
UXPS         GHKLEYESLRGKGIKALIAAPLRIFNAWEQARAIMKAYKPDVVLGMGGYVSGPGGLAAWS
UGAcS0.1     GHKLEYESLRGKGIKALIAAPLRIFNAWRQARAIMKAYKPDVVLGMGGYVSGPGGLAAWS
UGAcS0.2     GHKLEYESLRGKGIKALIAAPLRIFNAWRQARAIMKAYKPDVVLGMGGYVSGPGGLAAWS

UGAcS        LGIPVVLHEQNGIAGLTNKWLAKIATKVMQAFPGAFPNAEVVGNPVRTDVLALPLPRQRL
UXPS         LGIPVVLHERNGIAGLTNKWLAKIATKVMQAFPGAFPNAEVVGNPVRADVLALPLPRQRL
UGAcS0.1     LGIPVVLHEQNGIAGLTNKWLAKIATKVMQAFPGAFPNAEVVGNPVRTDVLALPLPQQRL
UGAcS0.2     LGIPVVLHEQNGIAGLTNKWLAKIATKVMQAFPGAFPNAEVVGNPVRADVLALPLPQQRL

UGAcS        AGREGPVRVLVVGGTTGARILNQTMPQVAAKLGDSVTIWHQSGKGSQQSVEQAYAEAGQP
UXPS         AGREGPVRVLVVGGSQGARILNQTMPQVAAKLGDSVTIWHQSGKGSQQSVEQAYAEAGQP
UGAcS0.1     AGREGPVRVLVVGGSQGARILNQTMPQVAAKLGDSVTIWHQSGKGSQQSVEQAYAEAGQP
UGAcS0.2     AGREGPVRVLVVGGSQGARILNQTMPQVAAKLGDSVTIWHQSGKGSQQSVEQAYAEAGQP

UGAcS        QHKVTGFIDDMAAAYAWADVVVCRSGALTVSEIAAAGLPALFVPFQHKDRQQYWNALPLE
UXPS         QHKVTGFIDDMAAAYAWADVVVCRSGALTVSEIAAAGLPALFVPFQHKDRQQYWNALPLE
UGAcS0.1     QHKVTEFIDDMAAAYAWADVVVCRSGALTVSEIAAAGLPALFVPFQHKDRQQYWNALPLE
UGAcS0.2     QHKVTGFIDDMAAAYAWADVVVCRSGALTVSEIAAAGLPALFVPFQHKDRQQYWNALPLE

UGAcS        KAGAAKIIEQPQLSVDAVANTLAGWSRETLLTMAERARAASIPDATERVANEVSRVARA
UXPS         KAGAAKIIEQPQLSVDAVANTLAGWSRETLLTMAERARAASIPDATERVANEVSRVARA
UGAcS0.1     KAGAAKIIEQPQLSVDAVANTLAGWSRETLLTMAERARAASIPDATERVANEVSRVARA
UGAcS0.2     KAGAAKIIEQPQLSVDAVANTLAGWSRETLLTMAERARAASIPDATERVANEVSRVARA
```

**Figure S4.** Sequence alignment of UGAcS with UXPS and two UGAcS early variants.

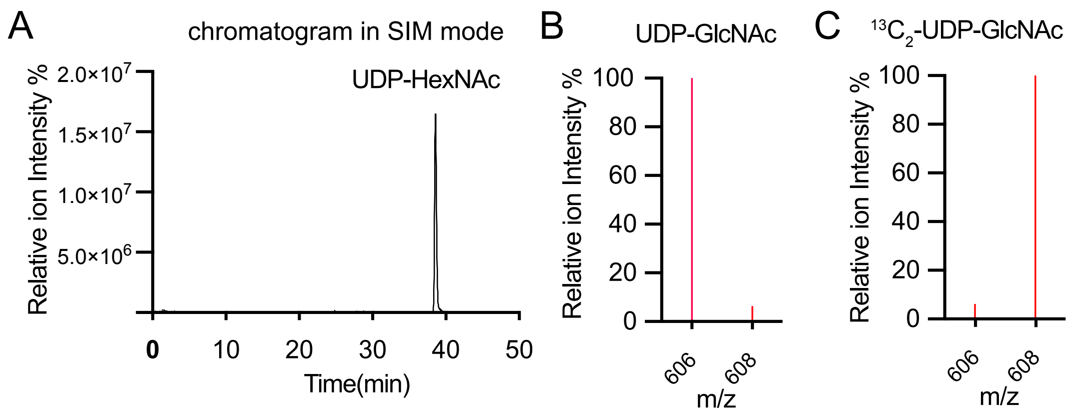


**Figure S5.** HILIC-MS analysis of relative UDP-HexNAc concentrations. (**A**) A representative chromatogram with the MS detector set in a selected ion monitoring (SIM) mode (m/z = 606 and 608), showing the elution at ~ 38.7 min. The peak represents the co-elution of UDP-GlcNAc and UDP-GalNAc (denoted as UDP-HexNAc). (**B**) MS analysis of an authentic UDP-GlcNAc sample, showing the dominant peak at m/z=606 and a minor isotopic peak at m/z=608. The relative abundance of the m/z=608 peak was 5.9% of the m/z=606 peak, close to the calculated theoretical value (5.4%). (**C**) MS analysis of $^{13}C_2$-UDP-GlcNAc, which was doped into the cell extracts as an internal standard.

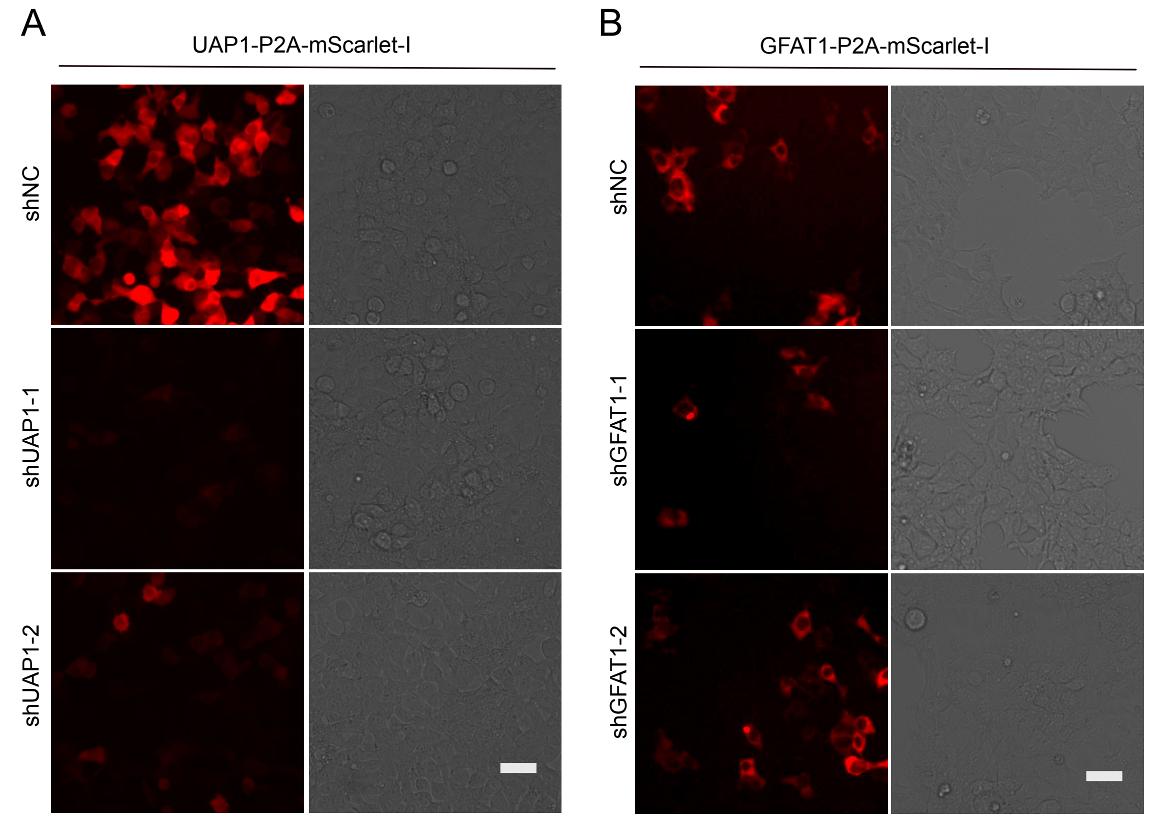


**Figure S6.** Evaluation of shRNAs for knocking down UAP or GFAT. (**A**) Images of HEK 293T cells expressing UAP1-P2A-mScarlet-I in the presence of shNC, shUAP-1, or shUAP-2 (Scale bar, 40 μm). (**B**) Images of HEK 293T cells expressing GFAT1-P2A-mScarlet-I in the presence of shNC, or shGFAT-1, or shGFAT-2 (Scale bar, 40 μm). The sequence of each shRNA is presented in the Methods section. shNC encodes a scrambled non-targeting sequence. shUAP-1 and shGFAT-1 were selected for further experiments because of their higher efficiencies than shUAP-2 or shGFAT-2.

### METHODS

#### Key Reagents and Materials

D-Glucosamine hydrochloride (GlcN), UDP-GlcNAc, UDP-GalNAc, UDP-Gal, UTP, CMP-Sia, GlcNAc-1-P, GDP-Fuc, GDP-Man, and UMP were purchased from MilliporeSigma (St. Louis, MO, USA). UDP was purchased from Cayman Chemical (Ann Arbor, Michigan, USA). 2-Deoxy-D-glucose (2-DG) was purchased from TCI America (Portland, Oregon, USA). $^{13}C_2$-UDP-GlcNAc was purchased from Omicron Biochemicals (South Bend, IN, USA). All oligos were ordered from Integrated DNA Technologies IDT (Coralville, Iowa, USA). pMD2.G (Addgene plasmid catalog no. 12259) and psPAX2 (Addgene plasmid catalog no. 12260) were gifts from Didier Trono (EPFL). pLKO.1 (Addgene plasmid catalog no. 10878) was a gift from David Root (Broad Institute).

#### Library Construction and Screening

The murG gene was amplified from *E. coli* DH10G and then inserted into a pBad vector via Gibson cloning. Next, the resultant pBad-murG plasmid was amplified with Phusion high-fidelity DNA polymerase and used for Gibson cloning along with a Circularly permuted green fluorescent proteins (cpGFP) fragment containing compatible ends. An NNK codon (N = A, T, G, or C; K = G or T) was introduced to each of the two ends of the cpGFP fragment. The library with two randomized residues at the two linkers was screened for UDP-GlcNAc responsiveness. Fluorescent colonies were selected for inoculation of cultures in 96-deep well plates with 1 mL 2×YT containing 0.01% L-arabinose and 100 μg/mL ampicillin in each well. The plates were shaken at 250 r.p.m and 37 °C for 5-8 h and then transferred to a 16 °C shaker for additional 48 h. Cells were lysed with 200 μL Bacterial Lysis Buffer (5 mg/mL octyl glucoside, 0.1 mg/mL chicken egg lysozyme, and 0.2 U/mL Benzonase in 20 mM Tris-HCl, pH 8) with 250 r.p.m shaking at 16 °C for 1 h. 600 μL of 50 mM Tris-HCl, pH 7.4 was added to each well of the 96-deep well plates. Next, the plates were centrifuged at 3000×g RCF for 10 min. 50 μL cell lysates from each well were transferred into two duplicate 96-well plates. 50 μL of 100 μM UDP-GlcNAc (prepared in 50 mM Tris-HCl, pH 7.4) was added to each well of the first 96-well plate, while 50 μL of 50 mM Tris-HCl, pH 7.4, as added to each well of the second 96-well plate. End-point fluorescence was measured on a BioTek Synergy Mx microplate reader. After the initial linker optimization, Gibson cloning with oligos containing degenerate codons was used to generate subsequent multisitedirected mutagenesis libraries, while error-prone PCR was used to create random mutagenesis libraries by following a previously described procedure.[1]

**Protein Purification**

A single colony of bacterial DH10G transformed with the UGAcS or UXPS gene in the pBAD vector was used to inoculate 2 mL 2×YT media supplemented with 100 μg/mL ampicillin. After overnight culture at 37 °C and 250 r.p.m, the cultures were used to inoculate 500 mL of 2×YT media with 0.02% L-arabinose and 100 μg/mL ampicillin. Cultures were incubated at 37 °C, 250 r.p.m for 2-5 h, and then incubated at 16 °C for additional 48 h. Cells were harvested via centrifugation at 5000×g RCF for 20 min. The cell pellet was resuspended in 30 mL buffer (50 mM sodium phosphate, 300 mM NaCl, and 10 mM imidazole, pH 7.4) freshly supplemented with 1 mM phenylmethylsulfonyl fluoride (PMSF). Cells were lysed with sonication. The cell lysate was centrifuged at 10,000×g RCF at 4 °C for 30 min using a JA-20 fixed angle rotor (Beckman Coulter). $His_6$-tagged proteins in the clarified lysate were purified with Ni-NTA agarose beads, and proteins eluted from the beads were subjected to size-exclusion chromatography through a HiLoad® 16/600 Superdex® 200 pg column (Cytiva) by using an elution buffer containing 50 mM Tris-HCl, pH 7.4. Purified proteins were further diluted with 50 mM Tris-HCl, pH 7.4 to a final concentration of ~ 100 nM and used for protein-based assays.

**Construction of Mammalian Expression Plasmids**

The UGAcS or UXPS gene in the pBAD vector was amplified and inserted into pcDNA3 between the Hind III and Apa I restriction sites. The UAP1 and GFAT1 genes were amplified from a human cDNA library constructed from the human embryonic kidney (HEK) 293T cells. The genes were genetically fused with P2A-mScarlet-I and inserted into pcDNA3 between the Hind III and Apa I restriction sites. pLKO.1 shRNA shuttle plasmids were constructed by following a previously described procedure.[2] The following RNAi target sequences were used:

shNC (scrambled non-targeting sequence)[3]: TTCTCCGAACGTGTCACGTAC

shGFAT-1 (TRC RNAi clone TRCN000 0075220)[4]: GCAGATACTTTGATGGGTCTT

shGFAT-2 (TRC RNAi clone TRCN0000075219)[4]: CGTCTTTCTATCCATCGAATT

shUAP-1 (TRC RNAi clone TRCN0000072371)[4]: GCCAATGATGTACCAATCCAA

shUAP-2 (TRC RNAi clone TRCN0000072372)[4]: CCAGACAAACCCAATGGAATA

#### Culture and Transfection of Mammalian Cells

HEK 293T cells were purchased from the American Type Culture Collection (ATCC) and cultured in Dulbecco's Modified Eagle Medium (DMEM) supplemented with 10% Fetal Bovine Serum (FBS), 100 U/mL penicillin, and 100 μg/mL streptomycin at 37 °C in a humidified incubator containing 5% $CO_2$. HEK 293T cells were plated on 12-mm glass coverslips in 24-well plates and grown to ~ 70% confluency. Cells in each well were transfected with 1 μg of plasmid DNA and 3 μg of linear, 25 kDa polyethyleneimine (PEI). Cell culture media were replaced 6-8 h post transfection. Cells were imaged ~ 18 h later.

Pancreatic β-cell line MIN6 cells were purchased from AddexBio and cultured in DMEM supplemented with 10% FBS, 0.0005% (v/v) β-mercaptoethanol, 20 mM 4-(2-hydroxyethyl)-1-piperazineethanesulfonic acid (HEPES; pH 7.4), and 1× Gibco GlutaMAX at 37 °C in a humidified incubator containing 5% $CO_2$. MIN6 cells were plated on 12-mm glass coverslips in 24-well plates and grown to ~ 70% confluency. Cells in each well were transfected with 1 μg of plasmid DNA and 3 μg of Lipofectamine 2000. Cell culture media were replaced 24 h post transfection. Cells were imaged ~ 18 h later.

#### Lentiviral Preparation and RNA Interference (RNAi)

The pLKO.1 shRNA shuttle plasmids mentioned above were used to transfect HEK 293T cells at ~ 70% confluency along with lentiviral packing and envelop expression plasmids psPAX2 and pMD2.G. Cell culture media were replaced 6-8 h post transfection and cells were cultured for additional 48 h. The cell cultures media contain lentiviruses, and were collected, filtered through 0.45 μm filters, and used without further purification. HEK 293T cells at ~ 70% confluence were infected with the crude lentiviruses for 3 days. Cells were cultured for another 5 days in the presence of 1 μg/mL puromycin. Cells were then transfected with the sensor plasmids as mentioned above for further experiments.

#### Fluorescence Microscopy

Fluorescence imaging was conducted on an inverted Leica DMi8 microscope equipped with a Leica TSC SPE-II spectral confocal module. Before imaging, transfected cells on a glass coverslip were transferred into a 35 mm glass-bottom dish containing 900 μL Leibovitz's L-15 Medium without phenol red (supplemented with 10 mM HEPES and 10% FBS). Cells were equilibrated at room temperature for ~ 2 h. Confocal images were acquired with a 40× oil objective (NA 1.3) under sequential excitation with 405 nm (~ 5% intensity) and 488 nm (~ 5% intensity) diode lasers. Emission was collected between 500 and 550 nm. For time-lapse imaging, the acquisition interval was set to be 2 min. Results presented in Figs. 3 and 5 were from fluorescence microscopy.

**Quantification of Cell Fluorescence with a Plate Reader**

Transfected HEK 293T cells in 24-well plates were subjected to various indicated treatments. Cells were rinsed three times with 200 μL Dulbecco's phosphate-buffered saline (DPBS), resuspended in 200 μL DPBS, and transferred into a 96-well plate. Fluorescence intensities were measured with a BioTek Synergy Mx microplate reader with excitation set at 400 nm and 488 nm, emission set at 520 nm, bandwidth set at 9 nm, and detector gain at 120. Fluorescence assay results presented in Figs. 2 and 4 were derived from plate reader measurements.

**Quantification of UDP-GlcNAc by HILIC-MS**

HEK 293T cells were seeded in 12-well cell culture plates and grown to near full confluency. Cells were treated with 10 mM 2-DG for 3 h (or untreated as the control). Cells were washed with 400 μL DPBS for 3 times and then scraped into 400 μL of cold DPBS. Cells were lysed with brief sonication. 1 mL cold ethanol and 5 μL of the internal standard (1 mM $^{13}C_2$-UDP-GlcNAc in water) were added to the cell lysate. Samples were next centrifuged at 12,000×g and 4°C for 20 min. The supernatants were collected, mixed with 4 mL dd$H_2O$, and lyophilized. Next, 20% acetonitrile in dd$H_2O$ was used to dissolve the lyophilized samples. Insolubles were removed by centrifugation at 12,000×g for 20 min. 50 μL of the supernatant of each sample was subjected to HILIC-MS analysis performed on a Waters LC-MS system equipped with a SeQuant ZIC-pHILIC PEEK-coated column (2.1×150 mm, 5 μm polymer; EMD Millipore, Billerica, MA), a Waters ZSpray™ multimode source, and a single quadrupole mass detector (Waters SQ Detector 2). Chromatographic conditions were as follows: flow rate was set at 0.19 mL/min, the mobile phase A consisted of 5mM ammonium acetate, pH 6.8 in water, the mobile phase B consisted of acetonitrile. The solvent program was 0-2 min, 80% B; 2-13 min, 80% to 40% B gradient; 13-14 min. 40% to 20% B gradient; 14-20 min, 20% B; 20-21 min, 20 to 0% B gradient; 21-28 min, 0% B; 28-30 min 0-80% B gradient; 30-50 min 80% B gradient. The mass detector was operated in an electrospray ionization (ESI) negative ion mode with selected ion monitoring (SIM) of 606 and 608 m/z ions. UDP-GlcNAc and UDP-GalNAc were co-eluted under the described chromatographic condition. The component is thus referred to as UDP-HexNAc.
